# Optimal organelle inheritance strategies under different changing environments and mutational pressures

**DOI:** 10.1101/2024.07.31.605980

**Authors:** Belén García-Pascual, Jan M. Nordbotten, Iain G. Johnston

## Abstract

Mitochondrial and chloroplast DNA (mtDNA and cpDNA) encode essential cellular apparatus. This organelle DNA (oDNA) exists at high copy number (ploidy) in eukaryotic cells, which must both mitigate mutational damage and allow adaptation to changing demands. Across eukaryotes, a range of inheritance strategies are used for oDNA, including uniparental and doubly uniparental inheritance (DUI), paternal leakage, recombination-mediated repair and gene conversion, and an effective “genetic bottleneck” imposed between generations. Here, we use modelling and simulation to investigate how these different strategies support the robustness and evolvability of oDNA populations under different challenges of mutation and changes in selection imposed by the environment. We find a general tradeoff between maintaining heterozygosity for flexible adaptation and supporting purifying selection against dysfunctional mutants. Different combinations of leakage and bottleneck size provide optimal resolutions to this tradeoff under different sets of challenges. The model explains many observed behaviours, including the appearance of non-minimal bottleneck sizes, a tradeoff between high ploidy for heterozygosity and repair and tight bottlenecks for segregation, and environmental dependence of the benefits of leakage and DUI. We connect different strategies observed across eukaryotes with the ecology of the organisms involved to explore support for the predictions of this theory.

## Introduction

Inheritance patterns of mitochondria and chloroplasts, and their internal organelle DNA (oDNA), are diverse across eukaryotes (Greiner et al., 2015; Munasinghe & Ågren, 2023). oDNA typically exists at high ploidy (hundreds or thousands of molecules per cell) and may be heteroplasmic, with an admixture of different oDNA variants within a cell (Johnston & Burgstaller, 2019; Van den Ameele et al., 2020; Wallace & Chalkia, 2013). How these multiple genomes are inherited between generations varies across lineages (Greiner et al., 2015; Munasinghe & Ågren, 2023; Veeraragavan et al., 2024). Some organisms follow the familiar human (and mammalian) pattern of strict maternal inheritance of mitochondrial DNA (mtDNA), with very limited or no “leakage” of oDNA from the father, and little or no recombination. Some bivalves show doubly uniparental inheritance (DUI), where either the mother or the father can contribute mtDNA to the offspring (Passamonti & Ghiselli, 2009; Zouros et al., 1992, 1994). Inheritance of mtDNA and chloroplast DNA (cpDNA, also called ptDNA for plastid DNA) in plants and algae is often maternal but sometimes paternal (Fauré et al., 1994; Greiner et al., 2015; Munasinghe & Ågren, 2023), and often involves leakage from the other parent (Breton & Stewart, 2015; McCauley, 2013). Plants show a dramatic amount of recombination in organelle DNA, involving templated repair and gene conversion processes (Broz et al., 2022, 2024; Khakhlova & Bock, 2006; Wu et al., 2020; Zwonitzer et al., 2024). Fungi and protists also use oDNA recombination (Barr et al., 2005), and different lineages show different modes of inheritance, including biparental inheritance (Birky, 2001), leakage (Wilson & Xu, 2012; Xu & Li, 2015) and even inheritance of mtDNA independently of either nuclear parent (three-parent offspring (Bloomfield et al., 2019)).

A particularly important aspect of organelle DNA inheritance is the “genetic bottleneck”, or the effective population size of organelle DNA molecules inherited by an offspring from its parent (Johnston, 2019b; Van den Ameele et al., 2020; Zhang et al., 2018). As inheritance has a random component, smaller bottlenecks introduce more variability in the oDNA profiles of offspring; larger bottlenecks mean a closer resemblance between siblings (and parent). A genetic bottleneck, manifest through a physical reduction in mtDNA copy number and parallel stochastic processes, is used to segregate mutational damage between offspring in animals (Johnston, 2019b). Plants also induce a genetic bottleneck, likely using a mechanism involving gene conversion and more limited physical reduction (Broz et al., 2022, 2024; Edwards et al., 2021). The inheritance of mtDNA in animals involves purifying selection (Fan et al., 2008; Lieber et al., 2019; Palozzi et al., 2022; Stewart et al., 2008), which is itself dependent on environmental cues (Latorre-Pellicer et al., 2019). Segregation of mutational load allows selection to act at the cellular or embryonic level, purifying population-wide oDNA populations (Edwards et al., 2021; Johnston, 2019b; Krakauer & Mira, 1999; Wallace & Chalkia, 2013; Zhang et al., 2018). MtDNA inheritance in animals, and presumably other taxa, thus involves an interplay of selection and segregation (Burgstaller et al., 2018; Johnston & Burgstaller, 2019; Veeraragavan et al., 2024).

Arguments have been put forward for the advantages and disadvantages of some of these strategies (Greiner et al., 2015; Munasinghe & Ågren, 2023). Quantitative models have been studied for the action of recombination (Albert et al., 1996; Atlan & Couvet, 1993; Lonsdale et al., 1997) and gene conversion (Edwards et al., 2021; Walsh, 1992), and the bottleneck and mutation-selection balance (Bergstrom & Pritchard, 1998; Roze et al., 2005). In addition to the above advantages of a genetic bottleneck and recombination, these arguments include uniparental modes limiting fitness costs associated with heteroplasmic populations (Christie et al., 2015; Hadjivasiliou et al., 2012; Lane, 2012; Sharpley et al., 2012) and promoting adaptative evolution (Christie & Beekman, 2017b, 2017a), and leakage helping circumvent “mother’s curse” and genomic conflicts (Beekman et al., 2014; Camus et al., 2012; Connallon et al., 2018; Cosmides & Tooby, 1981; Gemmell et al., 2004; Radzvilavicius et al., 2021). Theory has been advanced for how the evolution of an animal-like germline satisfies requirements for high-ploidy oDNA and mutational robustness (Colnaghi et al., 2021; Radzvilavicius et al., 2016), and how oDNA copy number shapes the relative influence of these different processes (Edwards et al., 2021; Roze et al., 2005).

Many of these modelling approaches have considered either no, or constant, selection. Here, we focus particularly on one class of proposed mechanisms supporting different modes of organelle DNA inheritance: adaptation to changing environments (Fraser & Kaern, 2009; Tufto, 2015). Previous work has argued that paternal leakage (Radzvilavicius & Johnston, 2021) and bottlenecks (in addition to their function in purifying selection) (Radzvilavicius & Johnston, 2022) can individually facilitate adaptation to changing environments; experimental work has shown that some environmental stressors influence plant organelle inheritance regimes (Chung et al., 2023). However, to our knowledge, a synergistic treatment of these strategies together (along with others like DUI and recombination-mediated repair) and considering both mutational hazard and environmental change, has yet to emerge.

Here, we attempt such a synergy through a stochastic modelling framework. We consider a portfolio of different organelle inheritance and maintenance strategies found across eukaryotes: bottlenecks, leakage, DUI, templated repair, gene conversion. We consider joint mutational and environmental change challenges to populations as they evolve and transmit organelle DNA between generations. Our main question is, if an organism is faced with changing environments and a mutational burden, what combination of strategies is optimal for organelle DNA inheritance? We show that different balances and magnitudes of external challenges effectively favour different weightings of genetic priorities in the evolving population, and identify the different strategies that are optimal with respect to these priorities. We also connect with behaviour observed across eukaryotic lineages and discuss the associated predictions that emerge from our model.

## Methods

### Model dynamics

The system consists of a population of N_pop_ organisms evolving over discrete generations (Fig. 1A). An organism is described by three values {A, B, M} describing the copy number of three alleles in its organelle DNA population: A and B (functional variants, which may have different fitness contributions in different environments), and M (dysfunctional variant). An organism is also assigned a sex, male or female, chosen so that exactly half the individuals in a generation are male and half are female. An organism’s fitness is given by a fitness function f(A, B) described below, with allele M never contributing to fitness. Time proceeds through discrete generations. A new organism is created as follows. A female mother and a male father are selected according to fitness through roulette wheel selection. Prior to mutation, an oDNA molecule is inherited with probability λ/2 from the father and with probability (1-λ)/2 from the mother:

**Figure 1.**
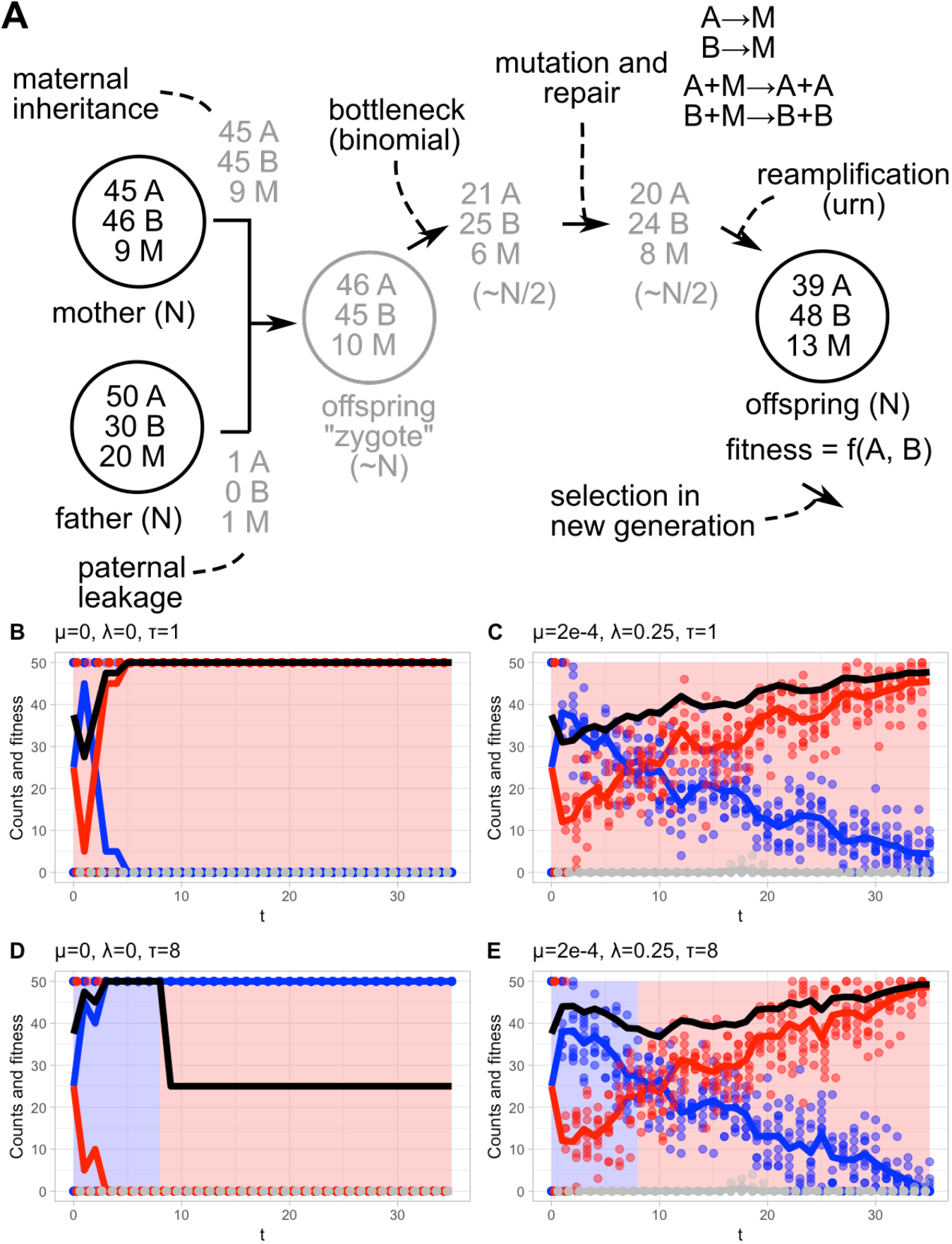
Model processes and illustrative behaviour. (A) *Constructing an o-spring*. Parents, randomly selected based on fitness, have {A, B, M} oDNA profiles (brackets give approximate copy number at each stage). These profiles are binomially sampled (potentially including leakage), combined, and subjected to a binomial bottleneck (Eqn. M1). Mutation (Eqns. M2-M3) and repair (Eqns. M4-M6) processes, if applicable, shape the oPspring’s oDNA profile. Reamplification (Eqn. M7) then produces an adult organism of the next generation. The process is repeated, sampling parents of the previous generation, until a new population of the same size is produced. Each is then assigned a fitness based on the fitness function and its oDNA profile, before selection for the next generation. (B-E) *Example model behaviours in di-erent conditions*. Points and traces give individual and population mean counts of A (blue), B (red), and M (grey) alleles. Black trace gives population mean fitness; background colour shows allele with selective advantage. (B) No mutation or leakage, constant environment: rapid fixation of the favoured allele (red). (C) Rare mutation and high leakage, constant environment: some mutational fluctuations and longer time to fix the favoured allele (not fixed by end of simulation). (D) No mutation or leakage, environmental change in generation 8: rapid fixation of initially favoured allele, no alternative allele retained, so later fitness drops. (E) Rare mutation and high leakage with the same changing environment: leakage preserves heterozygosity, allowing the system to adapt after environmental change.

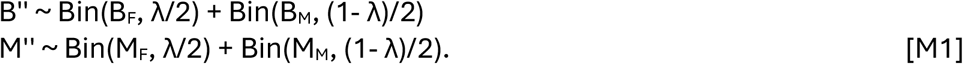

Asymmetric mutations are then applied with rate μ per molecule:

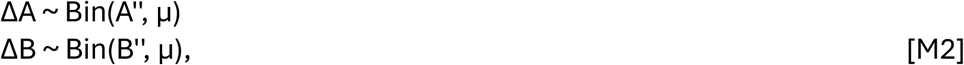

leading to oDNA profile

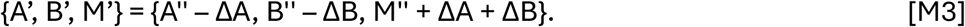

To simulate intracellular templated repair with rate ρ, we choose a number of mutants in an organism to undergo templated repair based on a randomly-chosen non-mutant oDNA molecule in the cell in a generation. This models the processes A+M → A+A and B+M → B+B, so that repair occurs at higher rates if stoichiometrically more A and B are available (Fig. 1A, (Zwonitzer et al., 2024)):

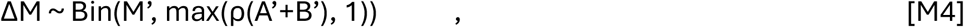

then assign them new identities

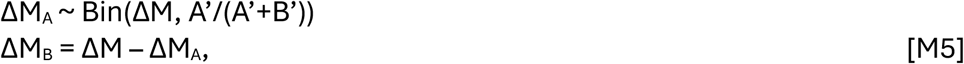

and update:

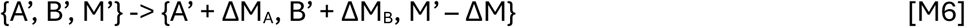

And finally, an urn-like model of random replication re-amplifies the oDNA population, where the process of randomly replicating a member of the current organelle profile (equal rates for all types) is repeated until the total copy number is N:

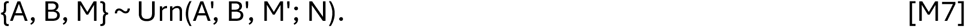

We therefore capture an effective “bottleneck” (sample size N/2) generating random variability upon inheritance, paternal leakage with rate λ, mutation rate μ, and selection according to a fitness function f(A, B).

To simulate doubly uniparental inheritance, every time a new individual is generated, the role of mother and father is flipped with probability 1/2 from standard (female mother, male father) to reversed (male mother, female father), then inheritance is simulated as above (with any nonzero leakage corresponding to inheritance from the father, which may be male (standard) or female (reversed)). Other model structures (deterministic reamplification, deterministic leakage, and heteroplasmic initial conditions) are demonstrated in Supp. Fig. 1.

### Environmental change

We model the environment of a population through the fitness function f(A, B). We will use two different environments:

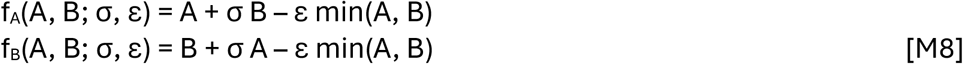

where σ <= 1 is the fitness contribution from the less-favoured allele in a given environment, and ε >= 0 is a heteroplasmy cost, penalizing mixed oDNA populations. Before considering selective differences we will use a simple case

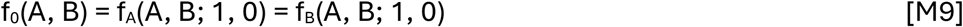

where both alleles contribute equally to fitness and there is no heteroplasmy penalty. We will also use

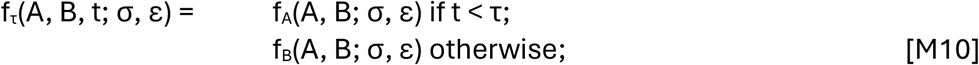

to capture environments that change from favouring allele A to favouring allele B at a timepoint τ. For initial conditions, we set half the males and half the females to be homoplasmic for A and half of each to be homoplasmic for B. We record population mean fitness after 500 generations. Some illustrative trajectories of the model (over a shorter timescale) are shown in Fig. 1B-C. The reader will notice similarities with, for example, the model of (Roze et al., 2005), but with some additional mechanisms, the inclusion of additional oDNA alleles, and a time-varying fitness function (Radzvilavicius & Johnston, 2022).

### Implementation

The model was implemented in custom-written C code. R (R Core Team, 2022) was used for visualization, with packages *reshape2* (Wickham, 2007) and *dplyr* (Wickham et al., 2023) used for data curation and *ggplot2* (Wickham, 2016), *metR* (Campitelli, 2021) and *ggpubr* (Kassambara, 2020) used for visualization. All code is publically available at https://github.com/StochasticBiology/odna-inheritance.

## Results

### Inheritance in constant environments

We first examined the “baseline” mutation-selection behaviour of our model, in the absence of changing environments. Previous work has derived expressions related to equilibrium fixation probabilities in a similar model (Roze et al., 2005), but to characterize our numerical model which does not necessarily reach an equilibrium state within the timeframe of simulation, we took a more empirical approach. We simulated the situation where σ = 0 and τ = 1 in Eqn. M10 (allele B has unit fitness and A and M have zero fitness). Example trajectories are shown in Fig. 1B-C. Across different values of paternal leakage and oDNA population size (bottleneck size), we found that the general expression

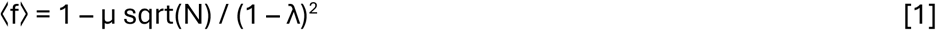

well described long-term mean fitness (Fig. 2; R^2^ = 0.96 across different organism population sizes, although larger leakage rates are less well correlated with the prediction).

**Figure 2.**
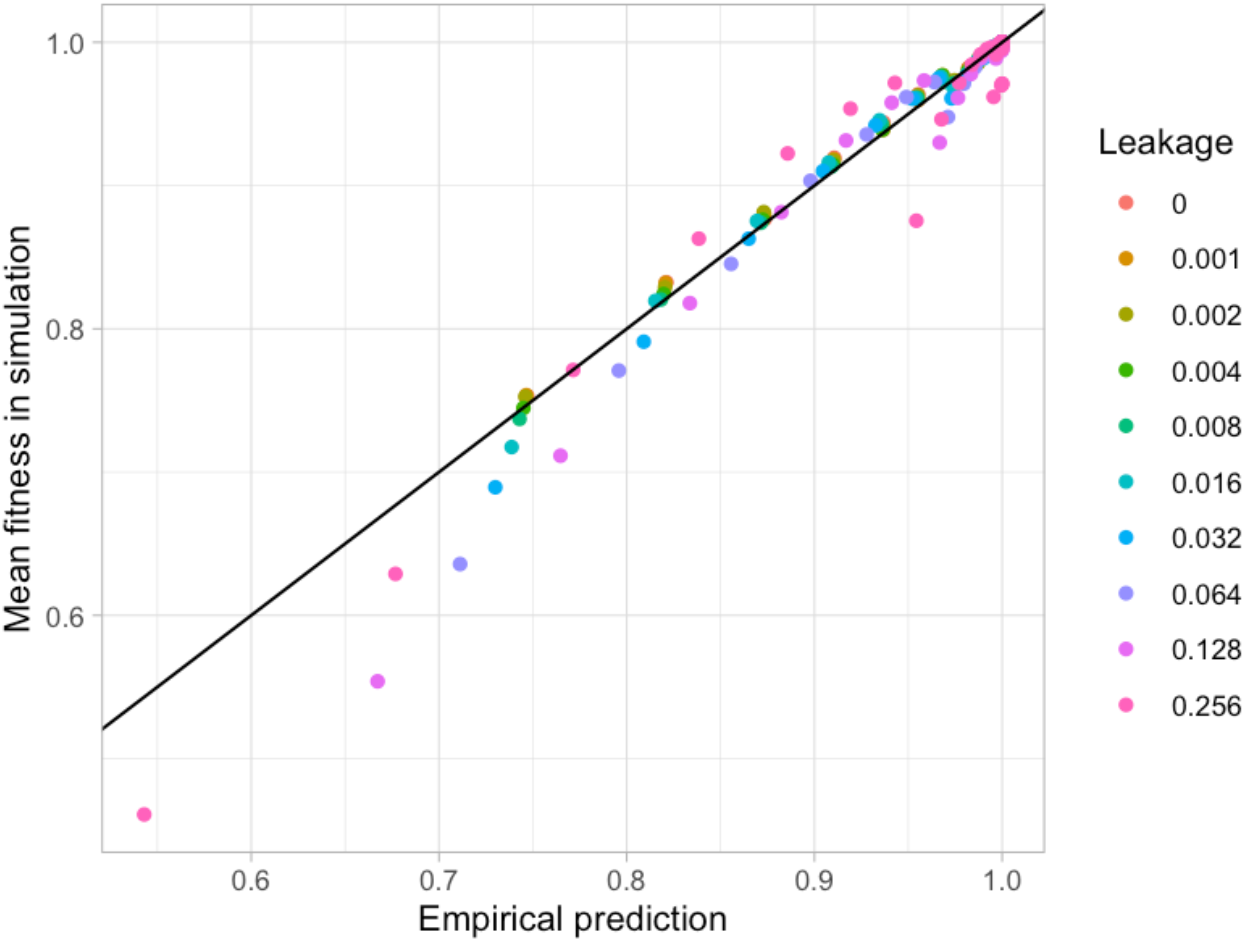
Empirical fit to constant environment behaviour. The horizontal axis gives an empirical prediction for mean fitness from the argument in the text; the vertical axis gives fitness observed in simulations. Colours give diPerent leakage rates; points within a colour correspond to diPerent μ and N values for that leakage. R^2^ = 0.96 for the predicted-observed link, but higher leakage values are less well correlated with prediction.

We do not aim to derive Eqn. 1 from first principles, but its form admits some intuitive analysis. First, in the case of no leakage and a single oDNA per organism, an organism’s fitness is determined entirely by the single allele it possesses, and we retrieve the expected result from classical population genetics that ⟨f⟩ = 1 – μ. The dependence on oDNA copy number N -- higher populations had marginally lower mean fitness for a given mutation rate -- is a consequence of a weaker “bottleneck” acting to generate selectable variability in mutant load between generations. In the infinite-N limit each offspring clonally inherits its mother’s mutant load; the variance of inherited mutant load h under binomial inheritance follows Var(h) ∼ 1/N, suggesting an interpretation of the sqrt(N) term -- the associated standard deviation -- as a measure of the selectable “spread” of values.

Nonzero leakage allows the bottleneck to be circumvented, supporting the maintenance of some mutant alleles that would otherwise be more efficiently purged by selection. The (1 – λ) empirical term in this sense resembles the (1 – μ) term above and in classical population genetics.

### Inheritance in changing environments

Having established this baseline behaviour, we turned to our main research question: which inheritance strategies are optimal under different regimes of environmental change? To this end, we simulated the population with a time-varying fitness function which changes from favouring allele A to favouring allele B after a given time window (example trajectories in Fig. 1D-E). We explored how the subsequent fitness of the population behaved under different time windows for environmental change and different mutation rates, as a function of paternal leakage rate and oDNA population size (Fig. 3).

**Figure 3.**
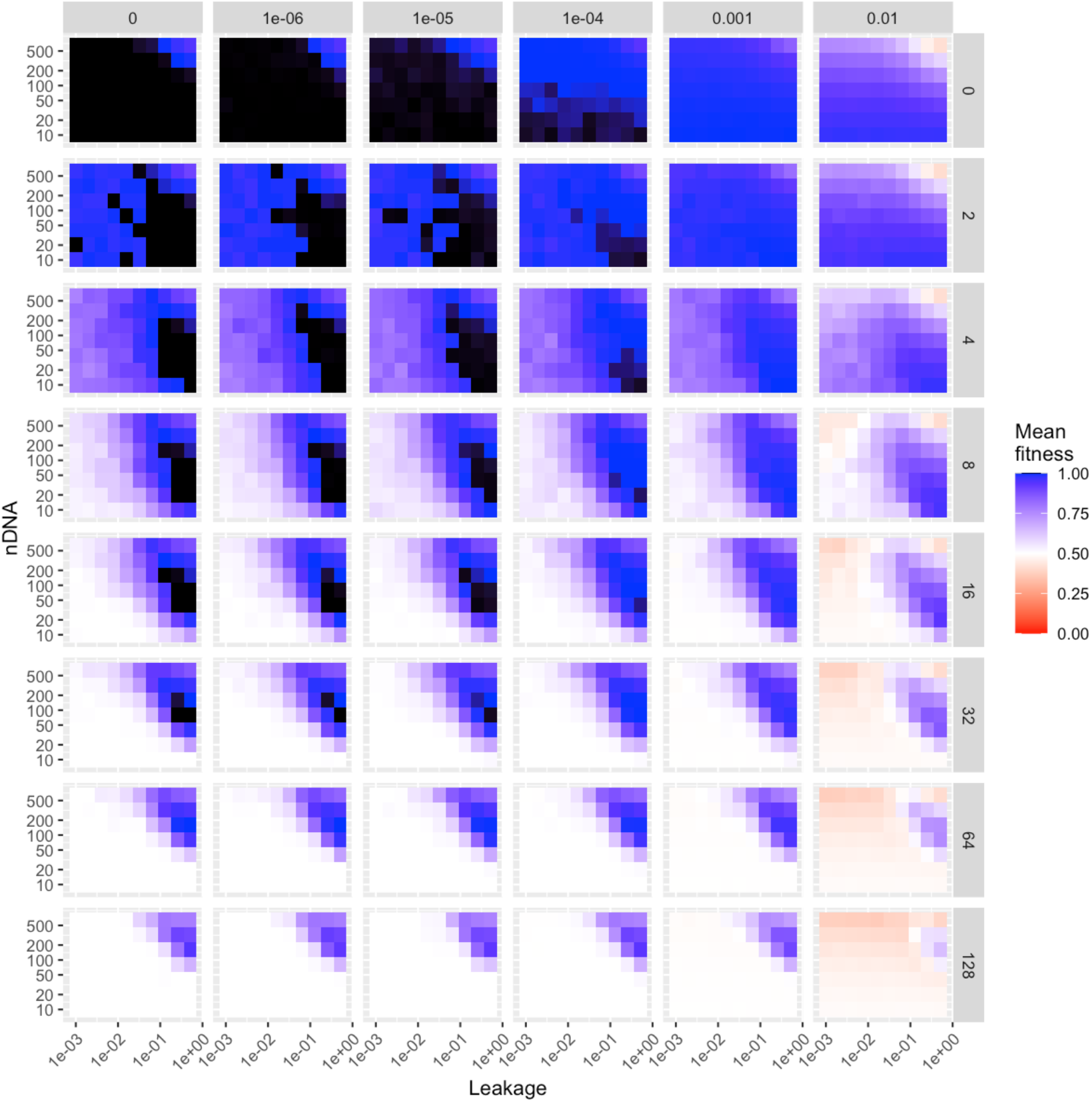
Fitness with di9erent inheritance strategies under di9erent challenges. Columns of the trellis give mutation rate, rows give time to environmental change event. In each panel, the horizontal axis is leakage rate and the vertical axis is oDNA copy number. Templated repair and DUI are both absent and there is no heteroplasmy penalty; organismal population size is 100. Supp. Fig. 2 shows comparable plots for alternative cases.

The case of zero environmental change (top row of Fig. 3) maps identically to the results in the previous section (Eqn. 1). For zero mutation rate, fitness is maximal in all cases except high N and high leakage, which increases the timescale necessary for complete adaptation beyond the scale of the simulation. Low N and low leakage reduce this timescale, leading to rapid fixation of the environment-matched allele. Low but nonzero mutation rates can be absorbed by the system, with the bottleneck acting to segregate mutants which can then be purged by cell-level selection. This negative selection is less effective for higher N (looser bottleneck), where the mutations that are not purged shift the system from perfect adaptation. At higher mutation rates (>10^−4^), even tight bottlenecks do not allow full purging of mutation, and some mutational entropy is always present in the system.

Any degree of environmental change changes this profile (lower rows of Fig. 3). Now, too-rapid fixation of the allele matched to the first environment becomes negative, as it removes from the gene pool the allele that is matched to the later environment. Hence, low N, low leakage behaviour is less optimal than higher leakage, which preserves heterozygosity for longer in the population and hence allows later fixation of the allele matching the second environment. Lower N values support the more rapid fixation of this second allele, with higher N values shifting from optimality as this fixation happens more slowly. For nonzero mutation, higher N values are further disfavoured as they diminish the power of the bottleneck to segregate mutation for subsequent purging. Low N, high leakage strategies are therefore favoured.

As the period of environmental change gets longer, the lowest N values become disfavoured, as they lead to too-rapid fixation of the first allele and loss of heterozygosity before the environmental change. There are then competing pressures on N even for zero mutation: too low and we lose heterozygosity, too high and we fail to adapt quickly to the later environment. For nonzero mutation, as before, high N values are further disfavoured due to a weak bottleneck’s inability to segregate damage. The priority of preserving heterozygosity leads to a coupling between leakage rate and N: high N / low leakage and low N / high leakage strategies allow the same heterozygosity, but high N / high leakage is challenged by the slow rate of later adaptation. As low N is valuable for purging mutations, we see low N / high leakage emerge as the optimal strategy under these conditions, with high N / low leakage a less optimal alternative, and other combinations substantially worse.

These trends are summarised in Fig. 4, where the optimal strategies for a given mutation rate and environmental change period are summarised. Optimal N decreases monotonically with mutation rate, reflecting the importance of the bottleneck for subsequent purging of mutation. For lower mutation rate, optimal N shows re-entrant behaviour with environmental change period: shorter and longer changes support higher N, while intermediate changes support lower N. This re-entrant observation is a result of averaging: the range of optimal N values is wider for shorter environmental change periods, and the average over all N that give maximal fitness is reported here.

**Figure 4.**
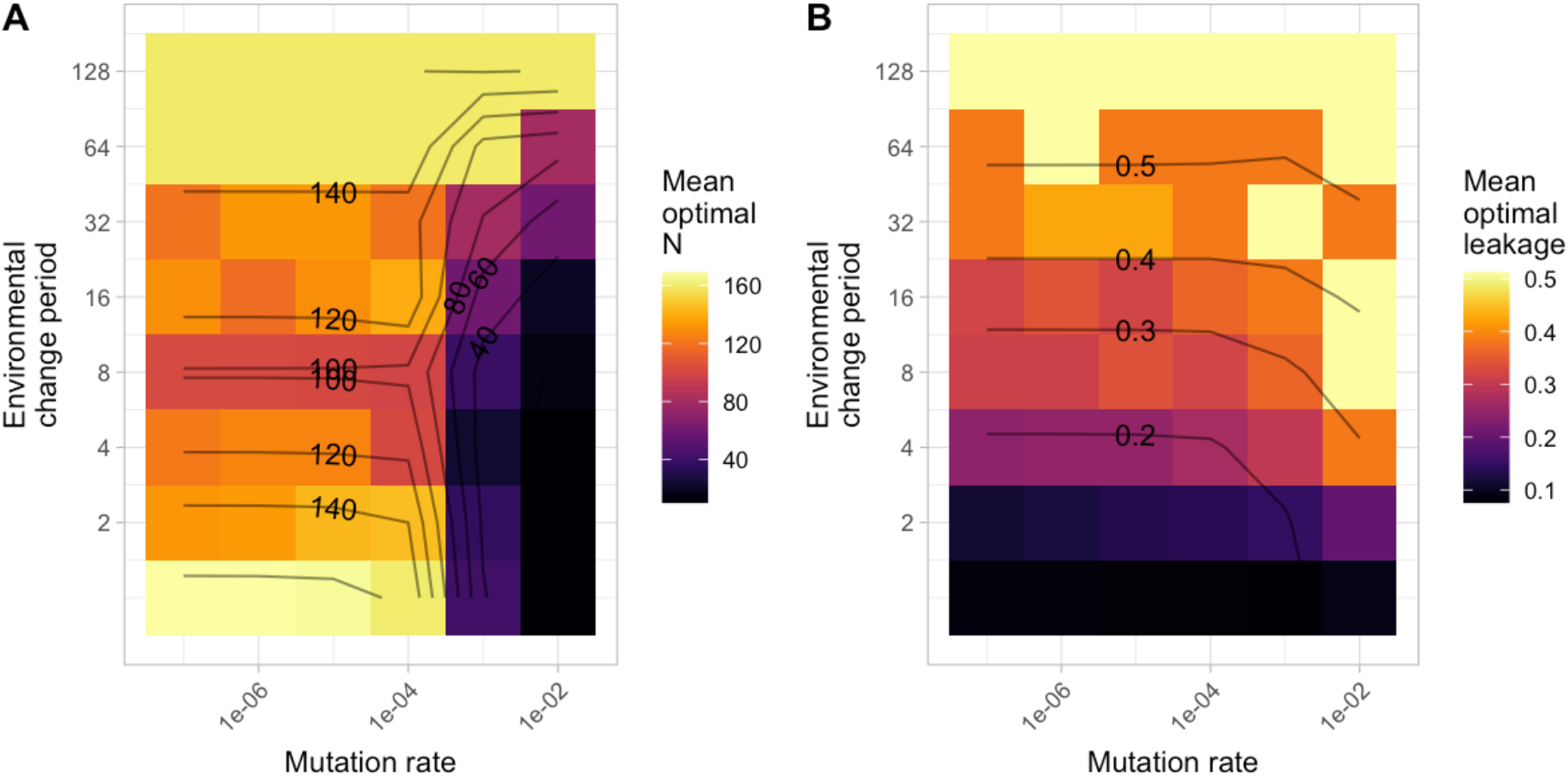
Optimal inheritance strategies under di9erent challenges. (A) Mean N value across strategies giving maximal fitness for a given mutation rate and environmental change frequency (zero on bottom row of pixels). An overall range of N from 1 to 512 was sampled. (B) Mean leakage rate across strategies giving maximal fitness. Contours are interpolated from a 2D LOESS fit across sampled values.

Optimal N becomes more constrained at low values for intermediate periods, and equally constrained to higher values at longer periods, so the trend is “flexible-lower-higher”.

The behaviour of optimal leakage is more straightforward, increasing strongly and monotonically environmental change period (supporting the maintenance of heterozygosity), and weakly and monotonically with mutation rate. This latter effect again comes from averaging: at low mutation rates there is a wider range of leakage values that support optimal behaviour (hence a lower average value), while this gets more constrained to higher values as mutation rate increases.

### Influence of different population structures and inheritance regimes

The analysis above is a basic, foundational model with fixed organismal population size, uniparental inheritance, no explicit heteroplasmy penalty for individuals, and no templated repair of mutational damage. We next asked how the profile of optimal strategies changed when we relaxed these assumptions (Fig. 5).

**Figure 5.**
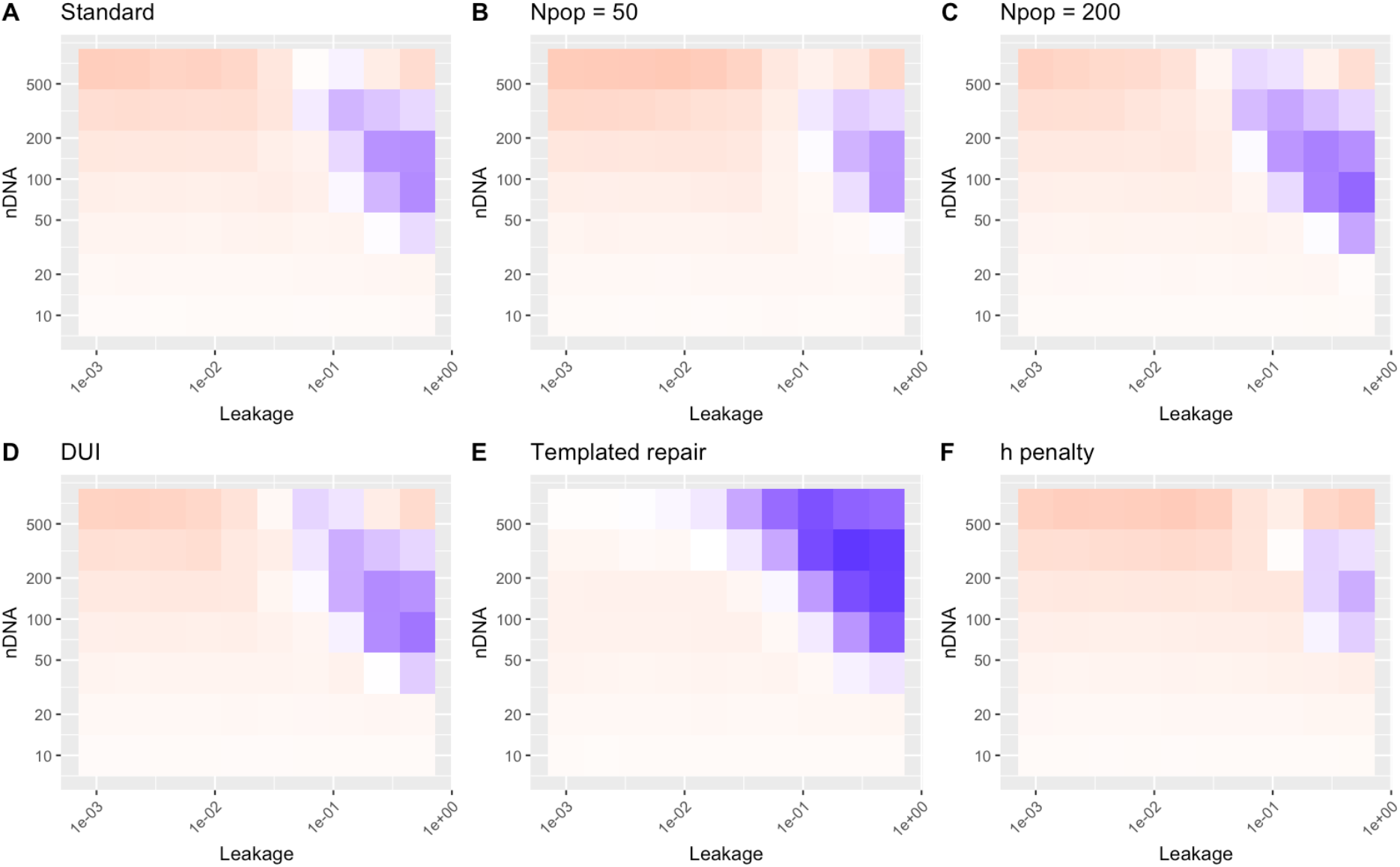
Influence of model variations. To demonstrate the influence of these effects, we focus on a particular instance of external challenge from Fig. 3 (environmental change period 64, mutation rate 0.01), though the trends we see here are reflected in other results (Supp. Fig. 2). Mean fitness is plotted as in Fig. 3, with the same colour scale (blue, high fitness; red, low fitness). (A) Default parameterisation (as in Fig. 3). (B) Smaller (N_pop_ = 50) and (C) larger (N_pop_ = 200) organismal population. (D) Doubly uniparental inheritance. (E) Templated repair with rate constant ρ = 10^−3^. (F) Heteroplasmy penalty ε = 0.25.

The influence of organismal population size and DUI is dominated by the strength of maintaining heterozygosity in a population. A smaller (0.5x) number of organisms has reduced capacity to retain heterozygosity and exhibits generally lower fitness (Fig. 5B), with high fitness constrained to a small region of strategy space and optimal behaviour slightly shifted to higher N (preserving heterozygosity taking priority over the bottleneck). Conversely, a larger (2x) number of organisms has an increased capacity to retain heterozygosity and performs better across a wider range of strategies (Fig. 5C), with optimal N shifted slightly downwards. Populations supporting DUI – in a simple model where mother or father is chosen randomly and uniformly as the oDNA source for an offspring -- resemble the behaviour of the (2x) larger population (Fig. 5D), somewhat intuitively, as the “pool” from which oDNAs can be inherited is now twice as large.

The introduction of templated repair (Fig. 5E) removes (to an extent dependent on parameterisation) the need for a bottleneck. This is because mutations can now be removed at the within-organism (intracellular) level, rather than relying on segregation across offspring followed by organism-level selection. As such, the pressure against high N due to weaker bottlenecking is largely removed. High N still challenges rapid fixation of the final allele, but this effect is offset both by the increased stoichiometric availability of wildtype oDNA to template repair of mutants (Zwonitzer et al., 2024) and by the increased capacity to preserve heterozygosity. With some amount of template repair, we thus see a substantial shift towards higher optimal N, and diminished re-entrant behaviour in N for a given leakage rate.

Finally, introducing an explicit fitness penalty for heteroplasmy provides a near-universal challenge to the system, and fitnesses are generally lower and optimal behaviour much more constrained (Fig. 5F). The maintenance of heterozygosity now comes at a fitness cost, promoting rapid fixation of the early allele and challenging the ability of the population to maintain and adapt the later allele. However, a small (depending on parameterisation) region of strategy space involving high leakage and intermediate bottleneck size is capable of retaining some fitness even in the challenging case demonstrated across Fig. 5. Here, the relative evolutionary advantage of heterozygosity for later adaptation balances the fitness cost to individuals of a heteroplasmic admixture. Evolution cannot of course “look ahead” to anticipate future change, but this result shows that a population adapted to leakage will not necessarily go extinct in this scenario when faced with an mtDNA admixture giving a strong heteroplasmy penalty.

## Discussion

The maintenance of organelle DNA in the face of environmental change and mutation rate involves several challenges with solutions that are sometimes competing. To preserve heterozygosity for adaptation to future challenges, rapid fixation of a single allele is not desirable – but rapid adaptation to current conditions is clearly positive. Preserving heterozygosity within individuals is easier with larger organelle population sizes – but this decreases the power of the bottleneck to segregate mutational damage for clearance. The relative importance and influence of these challenges depend on their parameterisation (biologically, the specific balances of rates, timescales, and other quantities), which varies tremendously across taxa and environments. But the direction of several of the challenges and strategies we study here can be summarised in Fig. 6.

**Figure 6.**
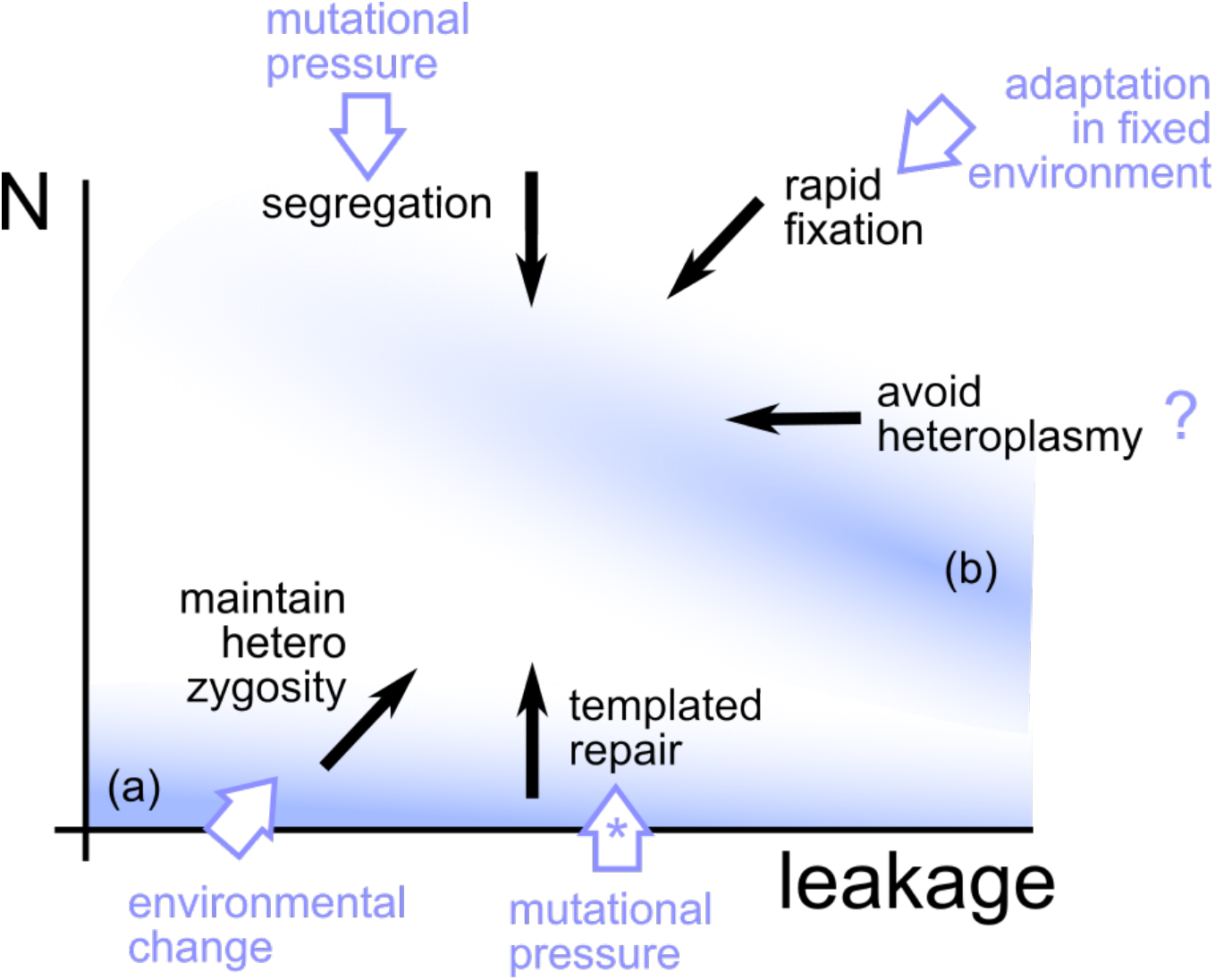
Summary of influences and strategies in our oDNA inheritance model. Black arrows give the directions of genetic priorities for a given system; the relative weightings of these ePects determine the optimal behaviour. White arrows illustrate how diPerent external pressures increase the relative importance of some priorities. Shaded blue regions illustrate (a) optimal region for static environments (Eqn. 1); (b) optimal region for changing environments. (?) the relative importance of avoiding heteroplasmy is hard to quantify on the same scale as the other values involved here, so this trend is presented semi-quantitatively. (*) templated repair is not supported by some lineages, in which case mutational pressure will typically favour lower N for bottleneck segregation and clearance.

What predictions can we extract from the model for biological interrogation? In agreement with previous work (Radzvilavicius & Johnston, 2021), lineages subject to more dramatic environmental change would be predicted to favour the preservation of heterozygosity through paternal leakage and DUI. Qualitatively, this agrees with, for example, the common observation of leakage in plants (Greiner et al., 2015) and DUI in bivalves (Passamonti & Ghiselli, 2009), both of which are sessile in dramatically changing environments. Environmental stress – specifically chilling stress – upregulates paternal leakage in plants (Chung et al., 2023). Heteroplasmy has long been observed in these lineages (McCauley, 2013; Pearl et al., 2009), as well as algae (Coyer et al., 2004) and fungi (Barr et al., 2005) often with similarly sessile lifestyles. Motile animals seem in general to exhibit fewer heteroplasmy-inducing and recombination strategies -- although, for example, some degree of leakage, heteroplasmy, and recently recombination-mediated repair have been observed in Drosophila (Klucnika et al., 2022; Nunes et al., 2013). Excitingly, recent advances in synthetically controlling inheritance strategies in plants (Chung et al., 2023) will afford the opportunity to test these hypotheses more directly.

What determines the optimal oDNA copy number in a given species? As illustrated in Fig. 6, several influences compete. Low N is beneficial for an animal-like bottleneck, although a bottleneck due to unbiased gene conversion can segregate damage independent of N (Edwards et al., 2021; Johnston et al., 2015; Walsh, 1992). High N is beneficial for templated repair (effectively, biased gene conversion): when mutant oDNAs are preferentially overwritten by wildtype oDNAs, having stoichiometrically more wildtype oDNAs is beneficial for mutant loads under 50% (Zwonitzer et al., 2024). High N is also beneficial for maintaining heterozygosity, as shown throughout this model. The particular N in a lineage then controls the balance between segregation due to an animal-like physical bottleneck (lower N) and repair of mutational damage (higher N) in that lineage, along with maintenance of heterozygosity (higher N), with gene conversion supporting segregation (independent of N) (Broz et al., 2022, 2024; Edwards et al., 2021). The balance of priorities between mutation rate, environmental change, and the efficiencies of intracellular repair versus cell-to-cell segregation of damage will then determine the optimal N in a given setting. Examples from plants indeed show tremendous variety in mtDNA copy number across species (Zwonitzer et al., 2024), with both repair and segregation facilitated by recombination machinery (Broz et al., 2022, 2024; Wu et al., 2020).

We have focused on genetic behaviour in our model, where a more fine-grained perspective would consider the physical behaviour both of organelles and of organisms. The physical dynamics of organelles influence both segregation and inheritance of oDNA (Aryaman et al., 2019; Edwards et al., 2021; Glastad & Johnston, 2022; Insalata et al., 2022; Tam et al., 2013, 2015), and the capacity for gene conversion and templated repair in recombining systems (Chustecki et al., 2021, 2022; Edwards et al., 2021; Giannakis, Chustecki, et al., 2022). Our single value N/2 for bottleneck size ignores the physical and genetic details of how this variability is generated, preventing a direct connection to the rates of the processes involved (Birky, 2001; Johnston & Jones, 2016). Expansion of this model to include the microscopic mechanisms of repair, segregation, and inheritance will be necessary to connect more quantitatively with true rates of these processes.

In parallel with the influence of changing environments on particular oDNA variants explored here, dynamic environments on the organismal, rather than generational, timescale have been argued to shape the structural evolution of the organelle genome (García-Pascual et al., 2022; Giannakis et al., 2024; Johnston, 2019a). Here, aligned with the co-location for redox regulation (CoRR) hypothesis (Allen, 2015), oDNA encoding of genes supports a rapid, individual-organelle regulatory response to changes in demand. A modelling approach ties this to changes in demand imposed by environmental conditions, arguing that strong environmental fluctuations favour oDNA encoding, while more stable environments favour nuclear encoding (García-Pascual et al., 2022; Giannakis, Arrowsmith, et al., 2022; Johnston & Williams, 2016). The interplay between beneficial oDNA retention patterns from that theory and beneficial oDNA inheritance strategies from this studies will be interesting to explore in future unifying work.

## Acknowledgements

This project has received funding from the European Research Council (ERC) under the European Union’s Horizon 2020 research and innovation programme (grant agreement no. 805046 (EvoConBiO) to I.G.J.).

## Supplementary Information

**Supplementary Figure 1.**
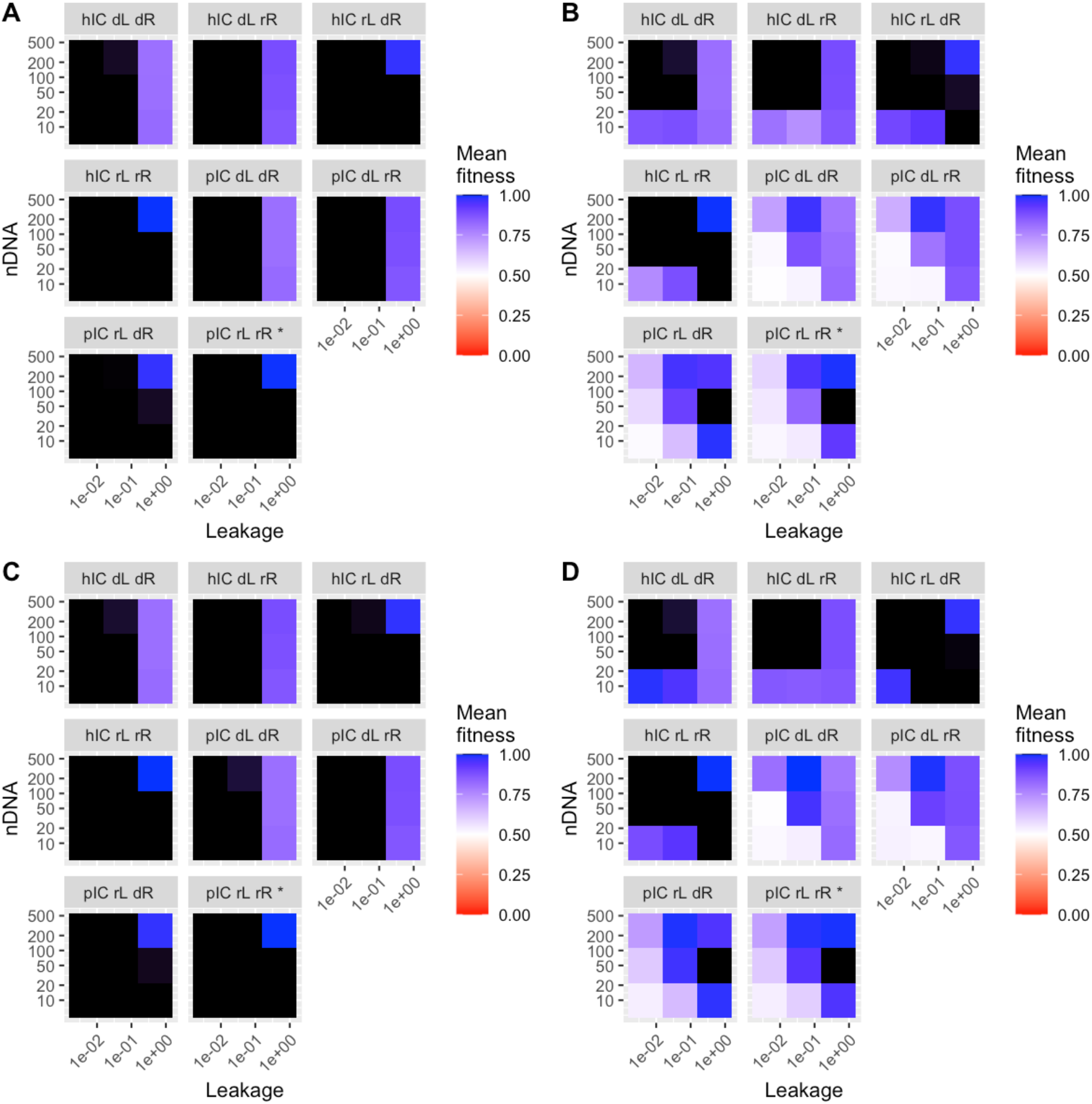
Fitness profiles under different models. (A) Static environment, no DUI; (B) Environmental change period τ = 10, no DUI; (C) Static environment, DUI, (D) Environmental change period τ = 10, DUI. Panels are labelled by model structure: h/p IC, heteroplasmic/pure (homoplasmic) initial conditions; d/r L, deterministic/random leakage; d/r R, deterministic/random reamplification. Asterisk shows model structure used in the main text. Heteroplasmic initial conditions dramatically change the dynamic profile, as leakage is no longer needed to create heteroplasmy. Deterministic leakage challenges adaptation; deterministic reamplification has a less pronounced effect (mimicking a slightly looser bottleneck).

**Supplementary Figure 2.**
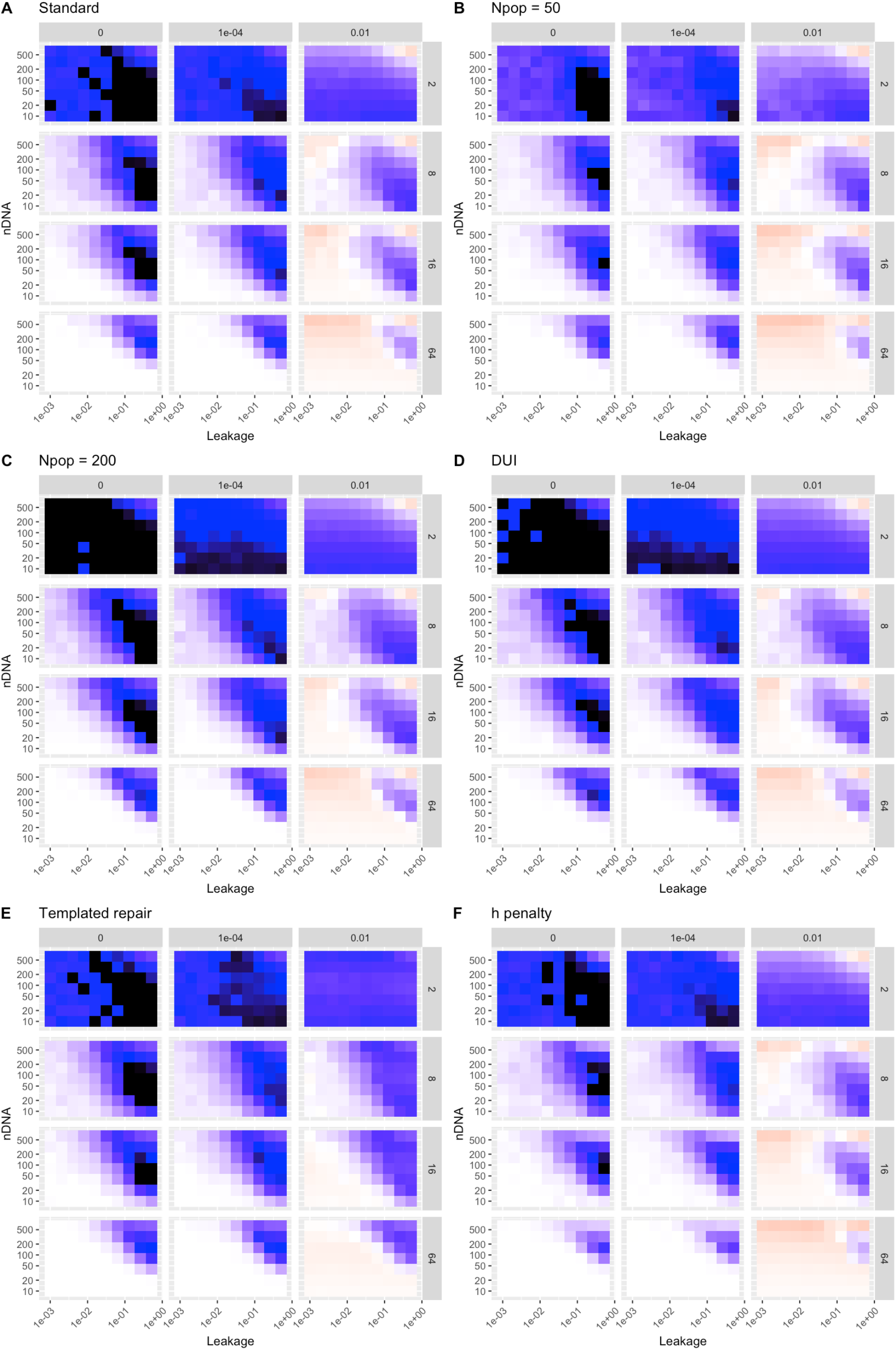
Strategy performance under model variations. Subsets of fitness plots with different inheritance strategies (horizontal axis, leakage; vertical axis, N) under different external challenges (mutation rate, columns; environmental change period, rows) for different models discussed in the main text and with the same colour scheme (black, highest fitness; blue, high fitness; red, low fitness).

## Notes

https://github.com/StochasticBiology/odna-inheritance

### Competing Interest Statement

The authors have declared no competing interest.

